# GM-CSF signaling on monocytes and monocyte-derived dendritic cells is required for effective pulmonary immunity to *Aspergillus fumigatus*

**DOI:** 10.64898/2026.09.22.753578

**Authors:** Kathleen A. M. Mills, Burkhard Becher, Tobias M. Hohl

**Affiliations:** Immunology and Microbial Pathogenesis Program, Weill Cornell Graduate School of Medical Sciences, New York, NY, USA; Institute of Experimental Immunology, University of Zurich, Zurich, Switzerland; Infectious Disease Service, Department of Medicine, Memorial Sloan Kettering Cancer Center, New York, NY, USA; Immuno-Oncology Program, Memorial Sloan Kettering Cancer Center, New York, NY, USA

**Author notes:** Center for Regenerative Medicine of Boston University and Boston Medical Center, Boston, MA, USA.

## Abstract

*Aspergillus fumigatus* is the most common cause of invasive aspergillosis (IA), an opportunistic pulmonary infection of immunocompromised patients that can be fatal, despite treatment with modern antifungal drugs. During infection, myeloid cells, such as neutrophils, monocytes, and monocyte-derived dendritic cells (Mo-DCs) are recruited to the lung and are critical for phagocytosing and killing *A. fumigatus* spores. Recent evidence has shown that intracellular crosstalk between these professional immune cells and other resident pulmonary cell types is required for an effective immune response to *A. fumigatus*, and that signaling via the cytokine granulocyte macrophage-colony stimulating factor (GM-CSF) is part of this intracellular communication. In this study, we sought to determine a role for GM-CSF signaling on monocytes and their progeny. We harnessed a mouse model of deletion of GM-CSF receptor β chain (encoded by *Csf2rb*) in CCR2-expressing cells. Using this model, we found that GM-CSF signaling is required on CCR2^+^ cells for killing of *A. fumigatus* and for host survival. This observation expands the importance of GM-CSF in host defense during *A. fumigatus* pulmonary infection and extends its role in pathways of intracellular crosstalk that mediate effective pulmonary antifungal immunity.

**IMPORTANCE:** *Aspergillus fumigatus* was recently designated a critical priority fungal pathogen by the World Health Organization due to its ability to cause fatal pulmonary disease and rapidly increasing acquired antifungal drug resistance. The mechanisms by which cells of the innate immune system cooperate to kill *A. fumigatus* spores in the lung are incompletely described and warrant further study so that these pathways may be manipulated in patients susceptible to infection. Here, we uncovered a role for the cytokine GM-CSF in coordinating crosstalk between lung epithelial cells and monocytes and monocyte-derived dendritic cells. GM-CSF signaling between these cells is required for fungal killing and host survival, expanding our understanding of the cell types involved in GM-CSF-mediated antifungal immunity.

## Introduction

*Aspergillus fumigatus* is the most common etiologic agent of invasive aspergillosis (IA), a deadly pulmonary fungal infection that can occur in immunocompromised patients (1, 2). Recruited myeloid cells, including neutrophils, CCR2^+^ inflammatory monocytes, monocyte-derived dendritic cells (Mo-DCs), and plasmacytoid dendritic cells, are essential for maintaining immunity to *A. fumigatus* as they phagocytose and kill inhaled fungal spores, termed conidia (1–5). Cellular crosstalk in homotypic and heterotypic interactions between immune and non-immune cells via cytokine mediators is an essential aspect of defense against *A. fumigatus* infection (4–7). We recently found that type 2 alveolar epithelial cells (AT2s) produce GM-CSF to license neutrophil killing of conidia following their recruitment into the lung (6). However, the effects of GM-CSF on the fungicidal properties of other cell types during *A. fumigatus* infection remain unclear. A limitation of studying GM-CSF is that mice lacking GM-CSF or either subunit of its receptor (GM-CSFRα or GM-CSFRβ) lack alveolar macrophages (AMs) as GM-CSF signaling is required for alveolar macrophage development and maintenance in the lung (8, 9).

Therefore, in this study, we sought to determine how GM-CSF affects the ability of monocytes and Mo-DCs to engulf and kill *A. fumigatus* conidia in an experimental model in which AMs are preserved. We found that GM-CSF signaling on CCR2^+^ cells (monocytes and their progeny) is required for optimal monocyte- and Mo-DC-dependent killing of *A. fumigatus* conidia and host survival after infection.

## Results

To determine whether GM-CSF signaling from AT2 cells is required for monocyte fungicidal activity, as it is in neutrophils, we used a tamoxifen-inducible mouse model of GM-CSF deletion in AT2 cells (*Sftpc-*cre^ERT2^ x *Csf2*^fl/fl^ mice, abbreviated *Csf2*^ΔSPC^) (6). We infected these mice and controls (either *Csf2*^ΔSPC^ mice given corn oil as a vehicle control or *Csf2*^fl/fl^ mice given tamoxifen), with <u>fl</u>uorescent *Aspergillus* <u>re</u>porter (FLARE) conidia (10) (Fig. 1A). FLARE conidia encode a constitutively expressed RFP and are surface labeled with AF633. Upon phagocytosis of live FLARE conidia, host leukocytes emit RFP and AF633 fluorescence, but upon fungal killing, RFP is rapidly degraded in the phagolysosome while AF633 fluorescence is maintained, resulting in AF633^+^RFP^-^ leukocytes. Leukocyte fungal uptake is quantified as the fraction of a given leukocyte population that is AF633^+^. To quantify *A. fumigatus* viability within leukocytes that have engulfed conidia, the frequency of RFP^+^ leukocytes is divided by the frequency of all AF633^+^ leukocytes. At 24 hours post-infection (hpi), we found that lung monocytes from *Csf2*^ΔSPC^ mice had comparable fungal uptake but increased fungal viability compared to monocytes from control mice (Fig. 1B&C). This finding indicates that AT2 cell-derived GM-CSF regulates the antifungal properties of lung monocytes.

**Fig. 1.**
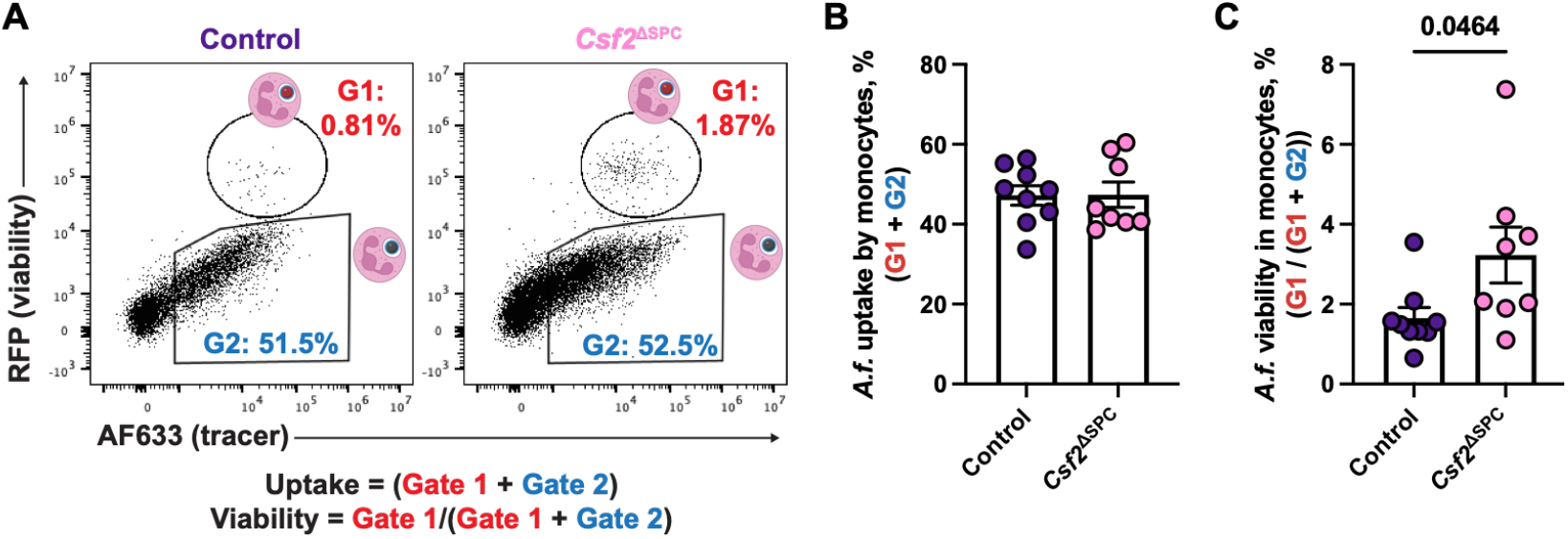
**(A)** Schematic of Fluorescent *Aspergillus* Reporter (FLARE) conidia and fluorescence emission after uptake and killing by lung monocytes from control or *Csf2*^ΔSPC^ mice. **(B)** Uptake of conidia by and **(C)** conidial viability in lung monocytes from control or *Csf2*^ΔSPC^ mice at 24 hpi with *A. fumigatus* conidia. Significance determined by Mann-Whitney test. Each symbol represents one mouse.

To determine whether this impairment in fungicidal activity by monocytes in the absence of GM-CSF production by AT2 cells is dependent on monocyte-intrinsic GM-CSF signaling, we crossed *Csf2rb*^fl/fl^ mice (11) to *Ccr2*-Cre mice (12). We confirmed that AMs were intact in this strain, as CCR2-depleter (*Ccr2*-DTR) mice were previously reported to have loss of some AMs after diphtheria toxin administration (3) and loss of AMs may be a confounding factor for anti-*Aspergillus* immunity (6). At both 24 and 48 hpi, *Csf2rb*^ΔCCR2^ mice had similar number of AMs compared to littermate controls (Fig. S1A&B). Additionally, *Csf2rb*^ΔCCR2^ mice had no defect in neutrophil accumulation in the lungs at 24 or 48 hpi (Fig. S1C&D), in contrast to *Csf2rb*^ΔMrp8^ mice in which *Csf2rb* is deleted in neutrophils (6).

To determine whether loss of *Csf2rb* expression in CCR2^+^ cells would impact mortality of *A. fumigatus*-infected mice, we quantified survival of these mice after infection. We found that, compared to littermate controls, *Csf2rb*^ΔCCR2^ mice had decreased survival rates, with 50% of mice dying by 3 days post-infection (Fig. 2A).

**Fig. 2.**
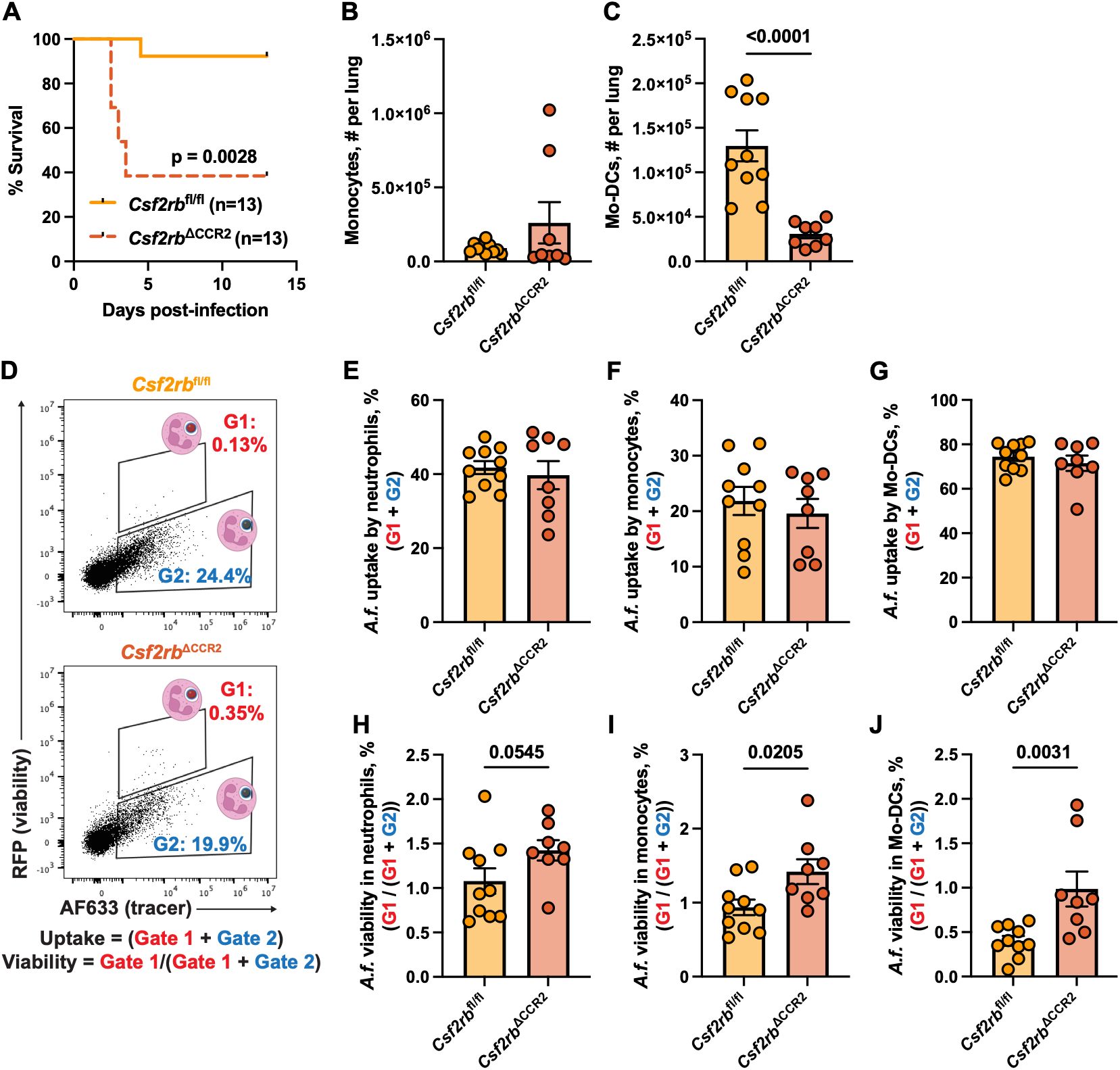
GM-CSF receptor signaling on CCR2^+^ cells is required for host survival, Mo-DC formation, and optimal fungal killing. **(A)** Survival of *Csf2rb*^fl/fl^ and *Csf2rb*^ΔCCR2^ mice after infection with *A. fumigatus* conidia. Significance determined by log-rank (Mantel-Cox) test. **(B)** Numbers of monocytes and **(C)** Mo-DCs per lung in *Csf2rb*^fl/fl^ and *Csf2rb*^ΔCCR2^ mice at 48 hpi. **(D)** Schematic of Fluorescent *Aspergillus* Reporter (FLARE) conidia and fluorescence emission after uptake and killing by leukocytes. **(E-G)** Uptake of conidia by and **(H-J)** conidial viability in lung neutrophils (E, H), monocytes (F, I), and Mo-DCs (G, J) at 48 hpi with *A. fumigatus* conidia. Significance determined by Mann-Whitney test. Each symbol represents one mouse. Results were pooled from two independent experiments.

Next, we wanted to determine whether *Csf2rb* deletion in CCR2^+^ cells affected their accumulation in the infected lung. There was no change in monocyte or Mo-DC numbers at 24 hpi (Fig. S1E&F). At 48 hpi, monocyte numbers were unchanged, however Mo-DC numbers were sharply curtailed in *Csf2rb*^ΔCCR2^ mice compared to littermate controls (Fig. 2B&C). These data are similar to those previously published by our group using whole-body *Csf2rb*^-/-^mice (13).

Finally, we wanted to investigate whether loss of *Csf2rb* on CCR2^+^ cells affected fungal uptake or killing using infection with FLARE conidia (Fig. 2D). At 24 hpi, we observed no differences in fungal uptake or killing by neutrophils or monocytes (Fig. S2A-D). There were too few Mo-DCs to analyze at this time point. At 48 hpi, fungal uptake was unchanged in neutrophils, monocytes, and Mo-DCs (Fig. 2E-G). However, at this time point, *A. fumigatus* viability was elevated in neutrophils, monocytes, and Mo-DCs from mice that lacked *Csf2rb* expression on CCR2^+^ cells (Fig. 2H-J).

## Discussion

In this study, we describe an essential role for GM-CSF signaling on CCR2^+^ cells (monocytes and Mo-DCs) in the anti-*A. fumigatus* immune response. AM, neutrophil, and monocyte numbers were unchanged during infection, though Mo-DC numbers were sharply curtailed, implying that GM-CSF signaling on monocytes likely regulates their differentiation into Mo-DCs. At 24 hpi, when neutrophil numbers peak in the lungs (5), fungal uptake and killing by neutrophils and monocytes is unimpaired. However at 48 hpi, when monocyte numbers peak in the lungs (5), fungal uptake by neutrophils, monocytes, and Mo-DCs is unaffected, but all three immune cell subsets exhibit an increase in fungal viability, i.e., impaired fungal killing, because a higher frequency of fungus-engaged cells contain live conidia compared to control mice with intact *Csf2rb* expression in CCR2^+^ cells.

These data suggest that GM-CSF signaling is required on CCR2^+^ cells for optimal intracellular fungal killing, and that monocyte or Mo-DC crosstalk with neutrophils downstream of GM-CSF signaling also regulates neutrophil-dependent fungal killing.

To date, the full scale of inflammatory mediators that facilitate monocyte-mediated fungal killing in the lung have been incompletely defined. Prior studies demonstrate that neutrophils license monocyte differentiation into Mo-DCs and subsequent fungal killing via production of interferon-gamma (IFN-γ) (7). In contrast to this IFN-γ related mechanism, the experiments with *Csf2rb*^ΔCCR2^ mice in this study indicate that direct GM-CSF receptor signaling on monocytes and Mo-DCs is required for optimal fungicidal activity.

Prior work in a mouse model of *Legionella pneumophila* lung infection found that GM-CSF signaling on monocytes increases their glycolytic capacity and promotes inflammatory cytokine production (14). Whether these same mechanisms are active in the context of *A. fumigatus* infection remains to be determined. In contrast to the beneficial effects of GM-CSF signaling on monocyte-dependent control of pulmonary *A. fumigatus* and *L. pneumophila* infection, CCR2^+^ monocyte ablation or global *Csf2* deficiency exacerbates murine pulmonary *Cryptococcus gattii* infection, highlighting pathogen-specific differences in monocyte- and GM-CSF-dependent requirements in host defense (15).

In sum, we show that GM-CSF derived from AT2 cells and signaling directly via the GM-CSFRβ chain on monocytes and Mo-DCs is required for optimal fungicidal activity of those cells and for host survival. Future work will seek to define the molecular mechanisms used by monocytes and Mo-DCs downstream of GM-CSF signaling.

## ACKNOWLEDGEMENTS

We thank all members of T.M.H.’s laboratory for insightful discussions. We gratefully acknowledge the support of the Research Animal Resource Center at MSKCC and Weill Cornell Medicine. The cartoon leukocytes included in FLARE flow cytometry plot diagrams were created in BioRender. This study was supported by NIH grants F31 AI167511 (to K.A.M.M.), P30 CA008748 (MSKCC, PI: C.M. Rudin), and R37 AI093808 (to T.M.H.). This study was also supported by the European Research Council (ERC) under the European Union’s Horizon 2020 research and innovation program (grant agreement no. 882424), the Swiss National Science Foundation (310030_170320, 310030_188450, and CRSII5_183478), and the Swiss Cancer League. The funders had no role in study design, data collection and analysis, decision to publish, or preparation of the manuscript.

## Notes

### Competing Interest Statement

The authors have declared no competing interest.

