## Supplemental materials for "GM-CSF signaling on monocytes and monocyte-derived dendritic cells is required for effective pulmonary immunity to *Aspergillus fumigatus*"

**MATERIALS AND METHODS**

*Mice*

*Sftpc*-CreERT2 mice (stock #028054) were purchased from The Jackson Laboratory. *Csf2*-floxed mice (1) and *Csf2rb*-floxed mice (2) were provided by Dr. Burkhard Becher (University of Zurich). *Ccr2*-Cre mice were developed by the Hohl Lab (3). The *Csf2rb*$^{\Delta CCR2}$ strain was generated by crossing these two strains together. Mice used in this study were 8-12 weeks old. Within experiments, mice were age- and sex-matched. Experiments were performed with both male and female mice. Mouse strains were bred and housed in the Research Animal Resource Center at MSKCC in individual ventilated cages under specific-pathogen free conditions. Animal experiments were conducted with approval of the MSKCC IACUC (protocol 13-07-008). Animal studies complied with all applicable provisions established by the Animal Welfare Act and the Public Health Services Policy on the Humane Care and Use of Laboratory Animals.

Aspergillus fumigatus *strains and murine infection model*

*Aspergillus fumigatus* strain CEA10 (provided by Robert Cramer, Dartmouth University) was used for all experiments. For experiments to analyze fungal uptake and killing using FLARE conidia, the CEA10-mRFP strain was used (4). *A. fumigatus* conidia were grown on glucose minimal medium slants for 4-7 days at 37°C prior to harvesting in PBS + 0.025% Tween-20 for experimental use. For FLARE experiments, 7.5 x $10^8$ conidia were incubated in 10 µg/mL EZ-Link™ Sulfo-NHS-LC-Biotin (ThermoFisher) in 1 mL of 50 mM $NaHCO_3$ buffer (pH 8.3) for 1-2 hrs at room temperature (or 16 hours at 4°C), washed with 1 mL of Tris-HCl (pH 8) buffer, incubated with 20 µg/mL Streptavidin, Alexa Fluor™ 633 conjugate (Molecular Probes) in PBS for 45 mins at room temperature, and resuspended in PBS + 0.025% Tween-20. To generate swollen heat-killed conidia, 5 x $10^6$/mL resting conidia were incubated for 16 hours in RPMI-1640 and 0.5 µg/mL voriconazole and then put in a heat block set to 100°C for 30 minutes. Mice were anesthetized by isoflurane inhalation and 3-6 x $10^7$ *A. fumigatus* conidia were instilled via the intratracheal route in 50 µL of PBS + 0.025% Tween-20. The standard inoculum is 3 x $10^7$ conidia (used for flow cytometry experiments). This dose is sublethal to most mouse strains on the B6 background, so we used 6 x $10^7$ conidia for survival experiments.

*Flow cytometry*

For analysis of immune cells, single cell suspensions of mouse lungs were generated by putting lungs in gentle MACS™ C tubes and mechanically homogenizing in 5 ml PBS using a gentle MACS™ Octo Dissociator (Miltenyi Biotec) in the absence of enzymes, then filtered through 100 μm filters. Red blood cells were lysed using RBC lysis buffer (Tonbo Biosciences), cells were blocked with anti-CD16/CD32, stained with fluorophore-conjugated antibodies, and analyzed on a Beckman Coulter Cytoflex LX. Single color controls for compensation were generated using lung cells or OneComp eBeads™ Compensation Beads. Experiments were analyzed with FlowJo version 10.8.1. Dead cells were excluded with DAPI or eBioscience™ Fixable Viability Dye eFluor™ 506 (ThermoFisher). Cells were gated as in Fig. S5 of reference (1). Alveolar macrophages are CD45+, CD11c+, Siglec-F+; neutrophils are CD45+, CD11b+, Ly6G+; monocytes are CD45+, CD11b+, Ly6G-, Ly6C+, MHCII-, CD11c-; Mo-DCs are CD45+, CD11b+, Ly6G-, Ly6C+, MHCII+, CD11c+.

*Analysis of fungal uptake and killing*

Mice were infected with FLARE conidia and lungs were harvested and prepared for flow cytometry. Uptake refers to the proportion of fungus-engaged cells, i.e., the fraction of a given cell subset that are $AF633^+$ (the sum of $RFP^+$ $AF633^+$ and $RFP^-$ $AF633^+$ cells). Viability refers to the proportion of fungus-engaged cells that contain live conidia, i.e., $RFP^+$ $AF633^+$ cells divided by the sum of $RFP^+$ $AF633^+$ and $RFP^-$ $AF633^+$ cells.

*Statistical analysis*

Statistical analysis was conducted with GraphPad Prism (versions 9 or 10). Tests used to determine significance are indicated in the figure legends. Data are displayed as mean ± SEM. Results were considered statistically significant at $p < 0.05$ (two-sided testing; alpha = 0.05). Data shown represent minimum two pooled experiments for each assay.

**Table S1. Key reagents.**

| REAGENT or RESOURCE | SOURCE | IDENTIFIER |
|---|---|---|
| **Antibodies** | | |
| Brilliant Violet 785™ anti-mouse CD45 Antibody (clone 30-F11) | BioLegend | Cat#103149 RRID: AB_2564590 |
| MHC Class II (I-A/I-E) Monoclonal Antibody (clone M5/114.15.2), Alexa Fluor™ 700, eBioscience™ | ThermoFisher | Cat#56-5321-82 RRID: AB_494009 |
| BD Pharmingen™ Purified Rat Anti-Mouse CD16/CD32 (Mouse BD Fc Block™) | BD Biosciences | Cat#553142 RRID: AB_394656 |
| BD Pharmingen™ FITC Rat Anti-Mouse Ly6C | BD Biosciences | Cat#553104 RRID: AB_394628 |
| BD OptiBuild™ BUV805 Rat Anti-CD11b | BD Biosciences | Cat#741934 RRID: AB_2871246 |
| BD Pharmingen™ PE-Cy™7 Hamster Anti-Mouse CD11c | BD Biosciences | Cat#558079 RRID: AB_647251 |
| BD Horizon™ BUV395 Rat Anti-Mouse Ly6G | BD Biosciences | Cat#563978 RRID: AB_2716852 |
| BD Horizon™ BV421 Rat Anti-Mouse Siglec-F | BD Biosciences | Cat#565934 RRID: AB_2722581 |
| Alexa Fluor® 647 anti-mouse Ly-6G Antibody | BioLegend | Cat#127609 RRID: AB_1134162 |
| BD Horizon™ BUV395 Rat Anti-CD11b | BD Biosciences | Cat#563553 RRID: AB_2738276 |
| **Chemicals** | | |
| Molecular Probes™ Streptavidin, Alexa Fluor™ 633 conjugate | Fisher Scientific | Cat#S21375 |
| Thermo Scientific™ EZ-Link™ Sulfo-NHS-LC-Biotin | Fisher Scientific | Cat#PI21335 |
| Voriconazole | Pfizer | N/A |
| Tamoxifen | Millipore Sigma | Cat#T5648 |
| Tween-20 | Sigma | Cat#P9416 |
| Isoflurane | Covetrus | Cat#029405 |
| Corn oil | Sigma Aldrich | Cat#C8267 |
| eBioscience™ Fixable Viability Dye eFluor™ 506 | ThermoFisher | Cat#65-0866-14 |
| RBC lysis buffer | Tonbo Biosciences | TNB-4300 |
| **Experimental models: Mouse strains** | | |
| *Csf2*-floxed mice | Burkhard Becher (University of Zurich) | N/A |
| *Sftpc*-cre$^{ERT2}$ mice | The Jackson Laboratory | JAX: 028054 |
| *Csf2rb*-floxed mice | Burkhard Becher (University of Zurich) | N/A |
| *CCR2*-cre mice | Tobias Hohl (MSKCC) | N/A |
| **Fungal strains** | | |
| CEA10 | Robert Cramer lab | |
| CEA10-mRFP | Robert Cramer lab | |

| | | |
|---|---|---|
| **Software and algorithms** | | |
| FlowJo 10.8.1 | FlowJo, LLC | N/A |
| Prism9 and 10 | GraphPad | N/A |
| **Other** | | |
| PBS 10X-Dulbecco's Phosphate Buffered Saline Solution-Liquid | Irvine Scientific | Cat#9242-500ML |
| Fisher BioReagents™ Bovine Serum Albumin, Heat Shock Treated | Fisher Scientific | Cat#BP1600-100 |
| OneComp eBeads™ Compensation Beads | ThermoFisher | Cat# 01-1111-42 |

### SUPPLEMENTAL FIGURES

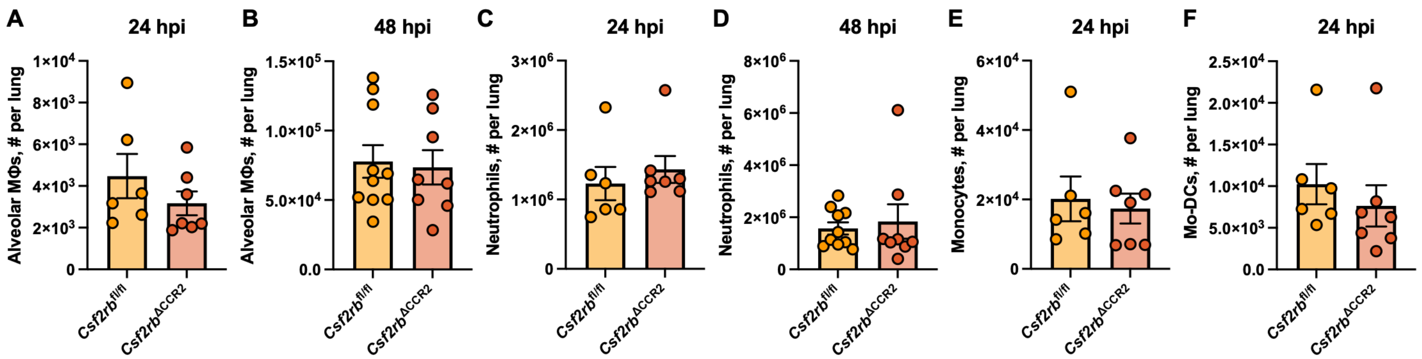


**Fig. S1. (A&B)** Numbers of alveolar macrophages, **(C&D)** neutrophils, **(E)** monocytes, and **(D)** Mo-DCs per lung in *Csf2rb*$^{fl/fl}$ and *Csf2rb*$^{\Delta CCR2}$ mice at indicated time points post infection. Significance determined by Mann-Whitney test. Each symbol represents one mouse.

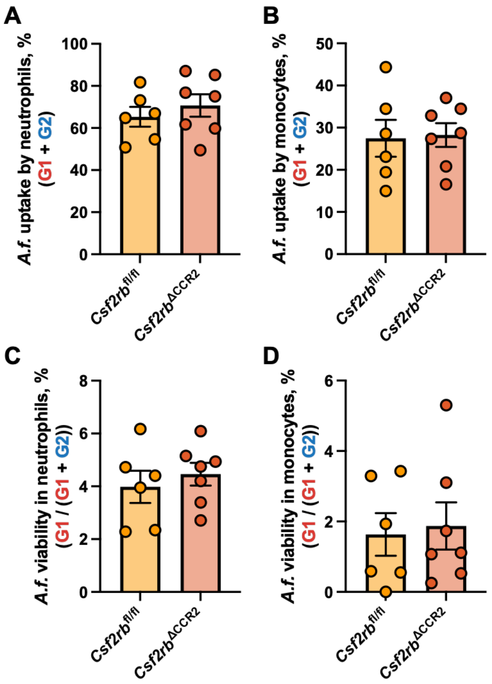


**Fig. S2. (A&B)** Uptake of conidia by and **(C&D)** conidial viability in lung neutrophils (A&C), and monocytes (B&D) from *Csf2rb*$^{fl/fl}$ or *Csf2rb*$^{\Delta CCR2}$ mice at 24 hpi with *A. fumigatus* conidia. Significance determined by Mann-Whitney test. Each symbol represents one mouse.
